# The power and pitfalls of Dirichlet-multinomial mixture models for ecological count data

**DOI:** 10.1101/045468

**Authors:** John D. O’Brien, Nicholas R. Record, Peter Countway

## Abstract

The Dirichlet-multinomial mixture model (DMM) and its extensions provide powerful new tools for interpreting the ecological dynamics underlying taxon abundance data. However, like many complex models, how effectively they capture the many features of empirical data is not well understood. In this work, we expand the DMM to an infinite mixture model (iDMM) and use posterior predictive distributions (PPDs) to explore the performance in three case studies, including two amplicon metagenomic time series. We avoid concentrating on fluctuations within individual taxa and instead focus on consortial-level dynamics, using straight-forward methods for visualizing this perspective. In each study, the iDMM appears to perform well in organizing the data as a framework for biological interpretation. Using the PPDs, we also observe several exceptions where the data appear to significantly depart from the model in ways that give useful ecological insight. We summarize the conclusions as a set of considerations for field researchers: problems with samples and taxa; relevant scales of ecological fluctuation; additional niches as outgroups; and possible violations of niche neutrality.

## Introduction

Metagenomic and amplicon sequencing techniques provide an unpredecented vantage to investigate the complexity of microbial ecologies, allowing researchers to simultaneously observe counts of thousands of species and will undoubtedly mold advances in human health, environmental science, and engineering (Riesenfeld et al., 2004; Steele and Streit, 2005; Tringe et al., 2005). The ability to observe whole ecosystems *in situ* has elevated microbiomes to a central position within ecology(Turnbaugh et al., 2007; Koenig et al., 2011; Aagaard et al., 2012; Caporaso et al., 2012; Delmont et al., 2012), These massive data sets present researchers with a new challenge: how to understand what they communicate about microbial ecological dynamics and, in turn, how this might inform ecology more broadly. This new project requires a dramatic shift in perspective, away from dynamics of small numbers of species to the interactions of thousands of them.

While metagenomic data adds a new urgency, taxon abundance data or ecological count data - collections where researchers estimate the number of observed individuals across a set of taxa within an ecosystem – is one the oldest forms of ecological measurement. Understanding the underlying phenomena governing how taxon abundance varies in time, space or along other environmental gradients is a foundational challenge within ecology (Bulmer, 1974; Etienne and Olff, 2005). A wide number of theoretical perspectives attempt to understand these variations, such as Lotka-Volterra and its descendants to models based on maximum entropy, food webs, trait-based dynamics, metacommunities, trophic cascades, mass and energy balance, and entropy production, among others (Record et al., 2013; Xiao et al., 2015; May, 2001; Hubbell, 2001; Allesina and Pascual, 2008; Lade et al., 2013; Leibold et al., 2004; Litchman and Klausmeier, 2008; Brown et al., 2004; Warren and Seifert, 2011; Petchey et al., 2008; Mougi and Kondoh, 2012; Alexander et al., 2012; Kleidon et al., 2010; Pace et al., 1999), While many approaches have compelling empirical or theoretical features, it is hard to argue that any have yet risen to provide a unified theoretical/empirical framework after the fashion of the coalescent model in population genetics (Rosenberg and Nordborg, 2002; Wakeley, 2009).

One of the most promising avenues to meet this challenge is the unified neutral theroy of biodiversity (UNTB), pioneered by Hubbell, that builds on the concept of neutral niches – collections of species that are ecologically equivalent – within a metacommunity structure (Hubbell, 2001; Harris et al., 2014). While frequently controversial, the UNTB has provided a versatile framework for understanding the structure of species abundance, particularly for species richness, dispersal, and island biogeography (Alonso and McKane, 2004; Dornelas et al., 2006; Rosindell et al., 2011). The crucial insight for the analysis of ecological count data is that, under suitable conditions, the Dirichlet-multinomial (DM) distribution is the sampling distribution of a neutral niche (Harris et al., 2014). If the count data for each sample are assumed to arise from a latent niche modeled by a DM distribution, the full metacommunity structure is given by a DM mixture model (DMM). (Holmes et al., 2012) gave the first practical inference for the DMM based on a variational Bayesian approach. Recent work by these authors shows how this framework can be broadened to an infinite-dimensional framework that can test for metacommunity neutrality (Harris et al., 2014). An independent extension uses a similar model for supervised feature detection (Shafiei et al., 2015).

In the rapidly evolving context of metagenomic statistical analysis, the DMM is only one of many approaches to make sense of observed count data. Unifrac, a phylogeny-based probability metric, as well as other distance based methods dominated early approaches and are still widely used (Lozupone and Knight, 2005; Chen et al., 2012). More recently, regression models based on single component DM and logistic normal distributions have given researchers a framework to understand how to associate significant shifts at the taxon level with environmental covariates while effectively accounting for structure of the overall count data (Chen and Li, 2013; Xia et al., 2013). An entirely distinct avenue has explored network-based analysis of correlations among taxa with increasing levels of statistical rigor (Liu et al., 2015; Biswas et al., 2015).

While appreciating the merits of these other approaches, here we focus on the DMM because of its statistical rigor, capactity for modeling extensions, and its link to the UNTB. For field researchers, the statistical truism that ‘all models are wrong, but some are useful,’ is naturally followed by the question ‘how useful are they?’ or, more pointedly, ‘how wrong are they?’ Our aim in this work is to give some answers to these questions for the DMM. Presented with a taxon abundance dataset, the DMM will always provide a means of organizing this data into distinct components; how well this organization can be relied upon is a necessary consideration for empirical ecologists. While the work of Harris et al. (2014) points to how the DMM may be re-embedded for general hypothesis testing, the integrated computational frameworks for statistical exploration common in other biological disciplines appear still far off (Drummond and Rambaut, 2007; Ronquist and Huelsenbeck, 2003). We emphasize that researchers can hold reasonable doubt about the verity of the UNTB, and still benefit from the DMM as a productive means of organizing and interpreting ecological count data (Ricklefs, 2006; McGill, 2003). This approach requires researchers to exercise due caution about the empirical features that may have be neglected or distorted by this analysis, such as communities dominated by a single species (e.g. algal blooms) or bias caused by a small number of samples.

To measure how well the model captures features of the data, we use posterior predictive distributions (PPD) to compare between the empirically observed data and the data that would be expected if the DMM were the correct underlying model (Gelman et al., 1996; Meng, 1994). PPDs are an important tool in Bayesian analysis, and provide a crucial means of assessing how adequately complex models are able to capture features of their underlying data sets. Here, we choose a set of test statistics intended to capture common statistical concerns relevant to ecological studies: within-sample taxa mean; across-sample mean for taxa; across-sample standard deviation for taxa; number of absent species within samples; and pairwise correlations among taxa.

Throughout the manuscript, we will refer to the components of the DMM as ecostates. This represents a departure from the common usage. (Holmes et al., 2012) use the term ‘ecotypes’ for these consortia, which is easily confused with its more common usage for a geographically-adapted subpopulation within a species. These components have also been labeled ‘metacommunities,’ though that term could also stand in for the collection of components (Nakatsu et al., 2015). Though ‘ecostates’ already has a referent within ecology (specifically, a leaf without a midvein), it is sufficiently distinct from the DMM and other closely-related topics that the meaning should be clear from context. We believe this phrasing also emphasizes a statistically pragmatic perspective about what these components represent: consistently inferred taxa consortia from a set of ecological samples, rather than niches or metacommunities per se.

In this investigation, we first outline a new model and inference scheme that generalizes the DMM to make use of a Dirichlet process mixture model (DPM). In it’s initial form, the DMM provides a single estimate, *K*, of the number of ecostates that best explain a set of count data (Holmes et al., 2012). However, in considering PPDs, a reasonable concern is that some model discrepancy arises purely from uncertainty in the choice of K rather than within the model itself. The DPM generalization of the DMM allows integration over all possible values of K and so directly accounts for this form of uncertainty. When necessary, we will refer to this generalization as the iDMM. We provide the scripts for this model under a Creative Commons License, written in the open-source R computing environment, available at the Github companion site for this paper: https://github.com/jacobian1980/ecostates.

We then apply the iDMM to three diverse time-series data sets from across ecology: well-known amplicon sequence count data collected from two individuals sampled over nearly 450 days at four bodily locations; amplicon sequence count data collected from the English channel approximately every two weeks over the year 2009-2010; and copepod abundance count data from continuous plankton recorders (CPR) collected during transects across the Gulf of Maine over the last fifty years. For each of these data sets, we use simple visualizations to emphasize how the iDMM reveals many previously identified features of the data, as well as pointing to some novel insights. The PPD analysis measures the overall agreement within the datasets, and we highlight problematic features and possible explanations within each. We conclude with a discussion including guidelines for field researchers using these methods for analysis.

## Data and Methods

### Data

We use three publicly available environmental time series as data sets. Each series was filtered for quality of metadata and the total number of counts per taxa. For the copepod time series, we filtered the data into three different sets based on taxon abundance. For the other two data sets a single set was used that screened out low abundance taxa. While each data set derives from a publically available archive, the specific data used in this paper have been archived at *figshare* with DOIs 3145321, 3145312, and 3145303. The filtered data sets are available on the companion website at:https://github.com/jacobian1980/ecostates.

### Human-associated bacterial abundance time series

To our knowledge, the most intensive amplicon-based, publicly available time series comes from 396 time points collected from four locations on two human individuals, frequently known by the title of the paper that described the data: ‘Moving Pictures of the Human Microbiome (MPHM) (Caporaso et al., 2011).’ This data was collected from the right and left palms, feces and saliva of a male and female subject over a total of 445 days from October 2008 until January 2010, for a total of 1967 samples. Illumina GAII sequencing targeted at V4 region of the 16S rRNA was applied to each sample and the resulting reads mapped to the Greengenes reference set, with the full protocol specified in (Caporaso et al., 2011). We downloaded the processed data and associated sample metadata from the Earth Microbiome Database on July 15, 2015. The data were filtered by taxon abundance for all taxa with more than 3100 counts across all samples, yielding 742 taxa in total.

### English channel bacterial abundance times series

The L4 station is a sampling location for the Western Observatory of the English Channel and the site of one of the first longitudinal metagenomic marine sequencing projects (Gilbert et al.,2009,2010; Caporaso et al., 2012). This collection builds on an extensive record of ecological and environmental sampling at this location dating back to 1903, with continuous plankton recording since 1998 (Record et al., 2010). This time series is the publicly available portion of a larger six-year series, and contains 68 samples gathered approximately 2 weeks apart from April 2009-April 2010. The large majority of samples were collected in pairs. Amplicon sequencing targeted the V4 hypervariable region of the 16S rRNA gene in every sample. We downloaded the raw data set and metadata including sampling times from the MG-RAST database on March 15, 2013 (Meyer et al., 2008).

We aligned the raw read data and screened for homopolymer artifacts using functions in the mothur software(Schloss et al., 2009). Reads were alligned to the SILVA reference alignment of 10,242 prokaryotic species (Release 102) (Quast et al., 2012). These were translated into species counts. At this stage, samples contained highly variable numbers of counts, from 2 to 267,529, with most samples near the median value of 33,041. We filtered these data into a single set for analysis, removing all taxa with fewer than 3 counts, leading to 994 taxa in total.

### Gulf of Maine time series copepod abundance data

Zooplankton are a crucial link in the ocean food web between the fish populations and microbial-scale trophic levels. For the past 70 years, continuous plankton recorders (CPR) monitored their abundance and diversity, via ships of opportunity that were transiting a region of interest. CPRs provide count data for identifiable copepod species at a depth of 5-10 meters below the ocean’s surface and have been used widely due to their low cost and ease of deployment. Here, we consider the copepod data collected via CPRs from 1964 to 2010 during transits across the Gulf of Maine (GoM), running from near Boston, Massachusetts to Yarmouth, Nova Scotia, available at the COPEPOD database (OBrien, 2005). While CPR data is among the most extensive marine ecological measurements, comparisons with metagenomic barcoding methodologies suggest that their total counts may be somewhat biased (Lindeque et al., 2013; Hirai et al., 2015).

The complete data contain 4799 samples, each enumerating the observed counts for a single CPR for 51 copepod species or genera. Each sample posssesses two metadata for the time of collection: position as longitude and latitude, and phytoplankton color index, a proxy measurement for the phytoplankton levels at the time of sampling. There is wide variablility in the number samples for each year, from ten in 1977 to 145 in 1989, with most measurements occurring between 1980 and 2000 (2456/4799). Each sample is located along an approximately one dimensional transect across the GoM. We filtered the samples to be located within 150 kilometers of the central line of this transect, that excludes 93 samples. An additional six samples were excluded due to ambiguous metadata.

## Methods

### Model

We employ an infinite dimensional generalization of the multinomial-Dirichlet mixture model of Holmes *et al.* (Holmes et al., 2012). This allows us to infer a posterior distribution that integrates over values of *K*, the number of components, removing this as a factor for aspects of the analysis. As in Holmes *et al.* we assume that each sample’s count data arises from a DM distribution. This distribution allows for additional dispersion relative to a strict multinomial distribution (Holmes et al., 2012). The model assumes that there are an unknown number of DM components (ecostates) underlying the data and that each sample comes from one of these components. Presuming the samples are otherwise exchangeable, a latent variable *c*_*i*_ augments each sample to assign it one of the ecostates. Supposing the DM paramers for an ecostate *k* are given by *A*_*k*_ = (*α*_1*k*_,⋯,*α*_*Mk*_), then, conditional upon *c*_*i*_ = *k*, the likelihood for a sample is given by

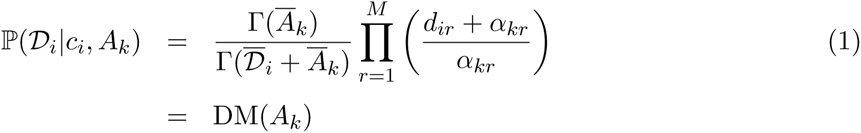

where Γ is the gamma function, 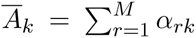 and 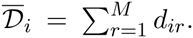. As the samples are assumed to be exchangeable, the full likelihood is then the product over all ℙ(*D*_*i*_|*c*_*i*_, *A*_*k*_):

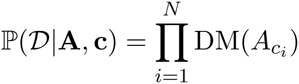

where **c** = (*c*_1_,⋯,*c*_*N*_). As in (Holmes et al., 2012), we adopt a Bayesian perspective on the problem. However, to remove *K* from consideration, we use a Dirichlet process mixture model (DPM), a nonparametric approach to specify the prior distribution on the parameters (Müller and Quintana, 2004; Teh et al., 2006). Following (Neal, 2000), the DPM can be formulated in this context as:

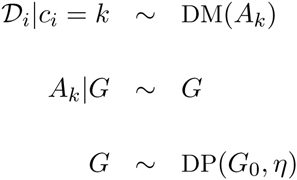

where ~ denotes ‘distributed as’, *G*_*0*_ is the base measure, *η* > 0 is a concentration parameter, and DP is a Dirichlet process. The base measure here is the Cartesian product of two independent distributions, with the first component an exponential distribution with mean one and the second a uniform Dirichlet distribution of length *M*.

### Inference

We use a Markov chain Monte Carlo (MCMC) methodology to approximate the posterior distribution of the model parameters, following the methods described in (Neal, 2000; Stein and Meng, 2013). All scripts were implemented in the **R** computing environment. We use two sets of Gibbs updates, one for the DPM parameters and one for the DM parameters. The DPM parameters (the latent variables) are drawn using a collapsed Gibbs sampler. At each iteration, the collapsed Gibbs update successively moves the sample’s latent variables to possibly new states, as specificed in Neal’s Algorithm 8 (Neal, 2000). For each sample *i*, we let *l* be the number of distinct culture labels *c*_*s*_ for *s* ≠ *i* and *h* = *l* + *m* where *m* is a parameter that allows for the assignment of a sample to a number of new components. We fix *m* = 3. If *c*_*i*_ = *c*_*s*_ for some *i* = *s*, then we sample values for that component. If *i* ≠ *s* for any *s*, then we set *C*_*i*_ = *l* + 1 and draw *m* new components from the base measure. Finally, we draw a new value for *c*_*i*_ according to:

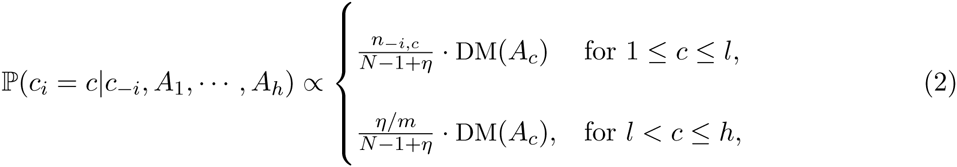

where *c*_‒*i*_ denotes all values of *c* except *c*_*i*_ and *n*_‒*i*_,_*c*_ is the number of values equal to *c* for *i* ≠ *s*. At each iteration, the set of *A*_*c*_’s is renumbered so all components have at least one associated sample.

The Gibbs steps to update the DM parameters come from recent work on how to generate efficient sampling from this distribution (Stein and Meng, 2013). The method relies on a data augmentation that separates the DM distribution into a Dirichlet distribution and a log-concave single parameter distribution. Since the Dirichlet is conjugate to the prior, it can be directly sampled. The last parameter, the dispersion parameters for the DM distribution, is sampled using a griddy Gibbs sampler with 1000 draws at each iteration (Ritter and Tanner, 1992).

For each dataset we ran three MCMC chains for 20,000 iterations. In each MCMC run, we applied several diagnostics to ensure convergence, including autocorrelation plots, Geweke’s diagnostic test, and estimating the effective sample size. We take a 10% of the chain as burn-in.

As an example, for one run of the full copepod data set, we realized minimal autocorrelation by thinning by 10. The Geweke statistic on the thinned chain was 2.047, consistent with reasonable convergence. The effective sample size was 423.78.

Scripts implementing the MCMC as well as visualizations routines are available on the companion website:https://github.com/jacobian1980/ecostates.

### Posterior predictive distribution

For a model specified by parameters Θ and data *D*, the posterior distrubution is the conditional distribution ℙ(Θ|*D*). The posterior predictive distribution for a new set of data *D̃* is generated by integrating the model likelihood of *D̃* over the posterior distribution:

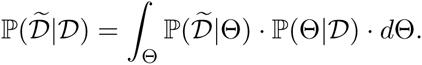

For complex models where this integral cannot be done analytically this distribution can be approximated by simulating a large number of datasets *D̃*_1_,⋯, *D̃*_*N*_ from ℙ(Θ|*D*). We then compare the observed data to these simulated sets through a test statistic *T*. Absent sufficient statistics, the appropriate choice of *T* is critical to reveal the empirical attributes of interest. These comparison are often summarized in terms of a posterior predictive *p*-value (PPP), the fraction of simulated test statistics more extreme than the empirically observed value:

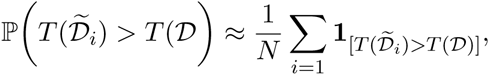

where 1_[.]_ is an indicator function.

To simulate the PPD, we used simulation routines in the R computing environment developed for the Human Microbiome Project (Turnbaugh et al., 2007). For each MCMC run, we simulated data in the following fashion. We thinned the Markov chain by 10 and trimmed it with a burn-in of 2,000 iterations. For each posterior sample and each data sample, we generated counts according to the corresponding DM distribution with the total number of counts equal to that observed in the data sample. We repeated this procedure 1,000 times, generating 50,000 simulated data sets per sample. We then calculated the PPP summaries for each taxa within each sample, the overal mean, overall standard deviation, number of zero-count taxa within a sample, and the pairwise correlations for the 15 most abundant and least abundant species using these simulated data sets. The PPD for each taxa within each sample was calculated by observing the fraction of simulated samples with more counts than the observed value. The mean, standard deviation, and pairwise correlation calculation were taken over all samples, providing metrics for each species. The zero calculation is taken over all species, provides a metric for each sample. All PPPs were then transformed by *p̃* = |0.5 – *p*|/0.5 so that values of either type of extremity yielded similar values.

### Visualizing the posterior distribution

Visualizing complex data is a crucial means to harness understanding from scientific inquiries. The iDMM effectively clusters the samples into a finite but variable number of components that reflect similarities in the count distributions across the samples. Most presentations using the DMM show taxon counts organized by these clusters to emphasize the specific shifts apparent across ecostates (Zhou et al., 2013; Charlson et al., 2012). This is a natural approach, particularly in a case-control study, where the researchers often seek to find the taxa that exhibit variation across experimental treatments. However, in the time series here, we prefer a presentation to highlight the ecostates’ relationship to each other and to time. This visualization is a simple organization of the posterior samples in a vein similar to the results of chromosome painting, so we term it ‘ecological painting’ (Hellenthal et al., 2008). We include the traditional, species-based presentations as Figures 2, 8,and 5 for comparison.

An ecological painting is simply a plot that colors samples according to their ecostate and orders the samples in time and space. In principle, other covariates may be used but these are natural first points of utilization. In the case of a finite mixture model, the number of components is fixed so colors can be assigned proportionally to the fraction of posterior samples in that state. In this case, we need to identify a homology of the ecostates across the posterior samples.

Unfortunately, there is no intrinsic aspect of the ecostates that render them identifiable across iterations. We deal with this problem by rendering the posterior into a finite mixture in the following way. First, we generate a coherence plot of the MCMC chain, showing the frequency of how often different samples fall within the same component. We also histogram the distribution of the number of components across the chain. From these we then determine a minimum number of components, *K′*, that reasonably capture the posterior sample. This value may not be the smallest number that the chain takes, and samples with components fewer that *K′* are excluded from the visualization. In our data sets, these iterations formed at most 2% of the chain. With *K′* in hand, for each iteration we find the largest *K′* components, and then enumerate all other components as *K′* + 1. This effectively reduces the chain to the finite case and so we apply the scheme propopsed by Stephens to minimize the Kullback-Leibler divergence between components across iterations (Stephens, 2000). Researchers may also be interested in how the ecostates relate to each other. To summarize this information, we present the expected frequency for the twenty most frequent taxa for each component.

## Results

We present the results for each data set, beginning with the biological interpretation of the ecological painting, followed by the indications the PPD gives about the model fit, and the correlation in ecostates between different level of taxa filtering. For each data set, we produce two figures – the painting and a summary of the PPD – for the purpose of comparison.

### Human-associated habitat bacterial time series

Figure 1 shows the ecological painting of the iDMM applied to the most heavily filtered set of the MPHM data. This presentation distills a number of key results from the original paper, rendering complex data amenable to visual analysis (Caporaso et al., 2011). Most salient is that the three niches (hand, feces, and tongue/saliva) are clearly separated by their inferred ecostates. Interestingly, the states assigned to tongue/saliva and hand habitats are shared across the two individuals, while, the feces habitat has a strong separation between them, consistent with the enterotypes hypothesis (Wu et al., 2011; Arumugam et al., 2011). For each habitat within each individual, the inferred ecostate exhibits strong temporal consistency. In particular, the female subject’s fecal samples exhibit near perfect consistency in ecostate across all time points. The ecostate painting shows the two hand pairs for both individuals exhibit strong correlation in their ecostates (*ρ* = 0.72, for maximum likelihood of the male subject’s samples), as observed in the original report.

**Figure 1:**
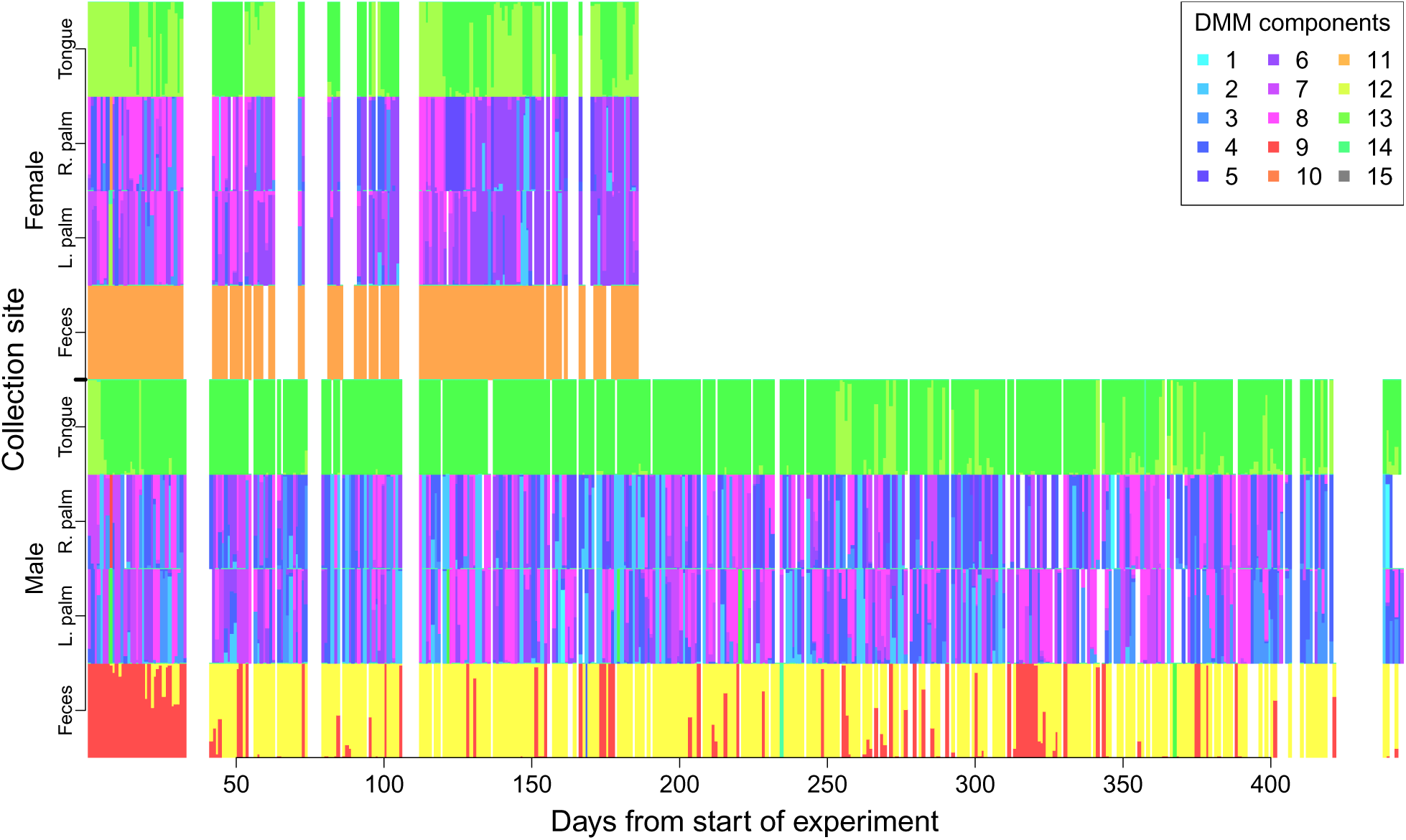
The ecological painting for the MPHM showing the mixture of ecostate for samples collected across two individuals (female above, male below) and the four collection sites over approximately 450 days. Each ecostate has a unique color noted in the legend.

The iDMM model also supports a more high-level analysis than a species-by-species comparison. For comparison, species distributions organized by ecostate can be seen in Figure 2. The strong temporal consistency in ecostate assignment across all habitats is modulated by regular oscillation within some of them. Within both individuals, the tongue/saliva habitat oscillates between two ecostates, with the duration of the oscillations lasting approximately one month for the male subject, and less than a week for the female subject. The painting also identifies a number of unusual samples, most obviously those colored green in the male hand and fecal samples. These may be mislabeled samples, or show unexpected associations in human microbiomes (between saliva and hand, for instance).

**Figure 2:**
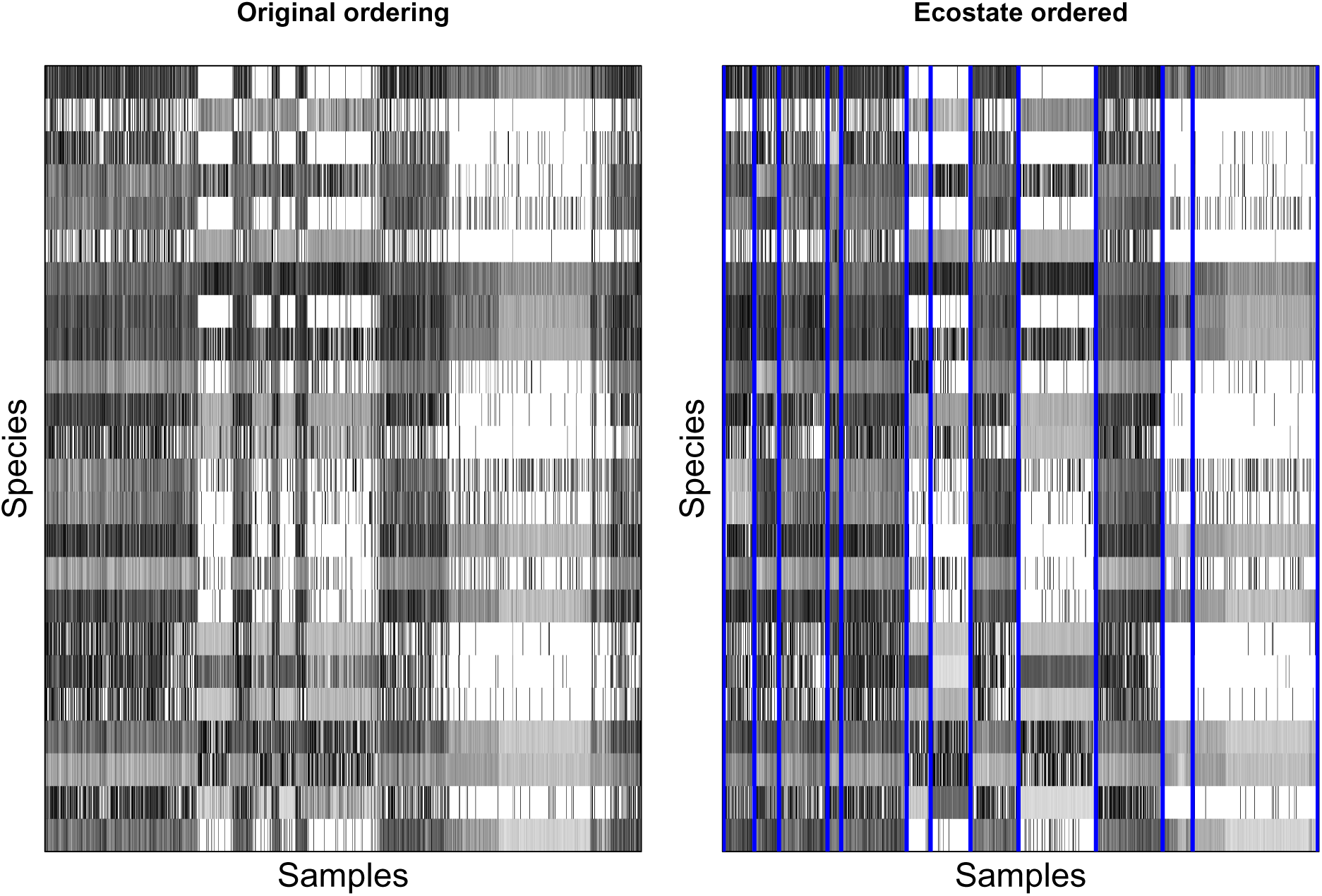
MPHM data set log-abundance arranged as in initial data set (left) and by maximum-likelihood ecostate (right).

Figure 3 summarizes the posterior predictive analysis for the data set. This indicates that the model effectively captures the mean and standard deviation in species counts and the frequency of zero counts in samples. The individual *p*-vales for each species across each sample (upper left panel) appear problematic but largely recapitulate the habitat structure, with reasonable *p*-values where particular species are common. The model struggles a bit to capture correlations across taxa, both at high and low levels of association, although they are almost always qualitatively correct (i.e. both are highly positive, or highly negative) if not quantitatively precise. Due to habitat structuring, the MPHM shows strong pairwise structure in correlation, with each pair of species showing high positive correlation (*ρ* > 0.5), low correlation (-0.5 < *ρ* < 0.5), or high negative correlation (*ρ* < –0.5). In nearly all cases, the model correctly estimates the category where a pair of species falls. However, the empirical fit provides reasonable *p*-values for the low-correlation cases. Even controlling for habitat structure (only considering pairwise correlation among species prevelant in the same habitat) this pattern persists. This indicates that the iDMM is useful for qualitatively capturing species correlations but does not provide precise quantitative estimates across disparate niches.

**Figure 3:**
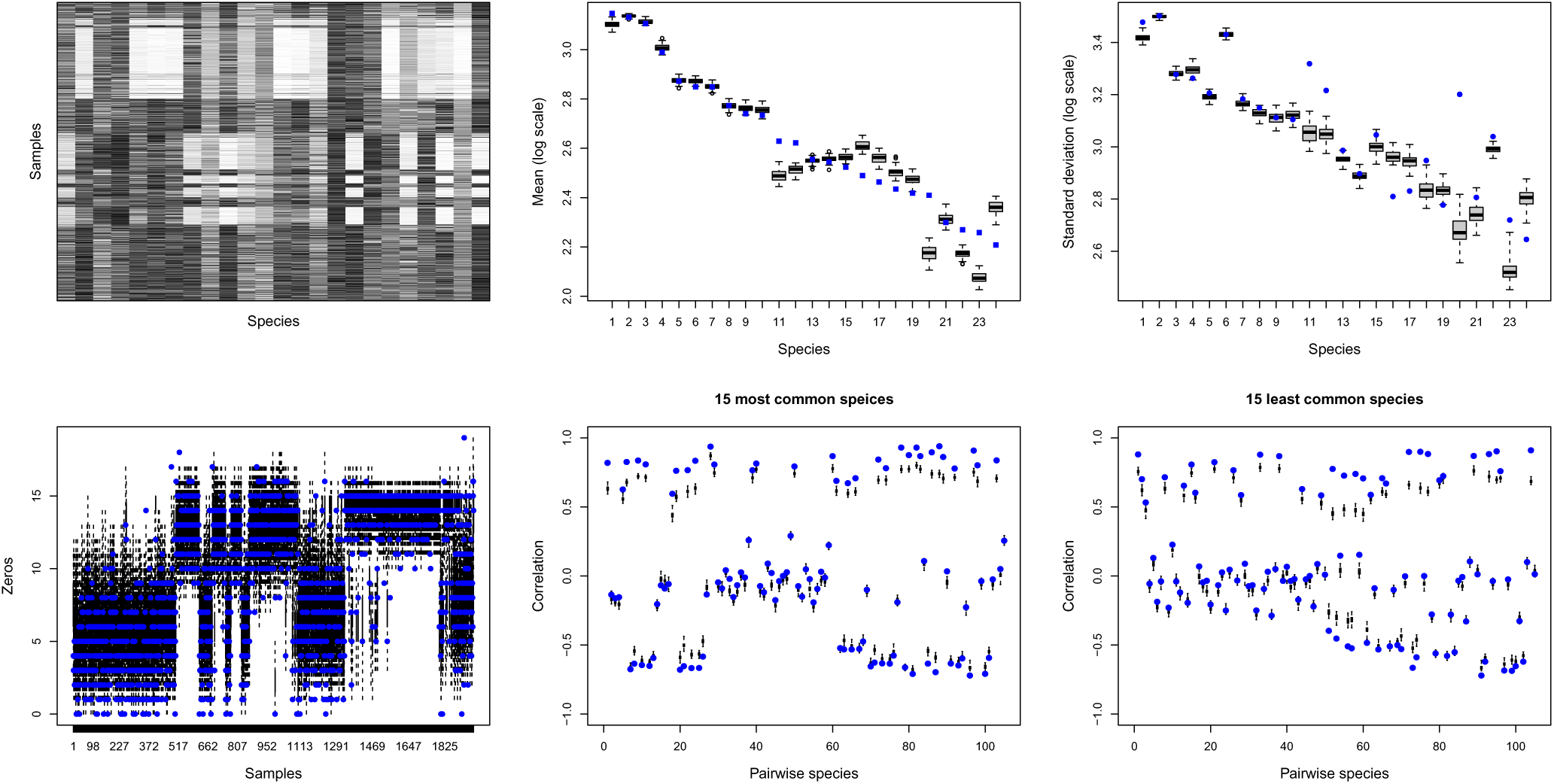
The PPD for the MPHM showing performance for each taxa within each sample (upper left), mean across all samples (upper middle), standard deviation across all samples (upper right), zeros within samples (lower left), pairwise correlation among abundant species (bottom middle) and infrequent species (bottom right). Species ordered by total abundance. Samples orderd by habitat. Upper left panel has extreme *p*-values in white. Boxplots show simulated values; blue dots show observed values.

### English channel bacterial time series

Figure 4 shows the ecological painting for the L4 time-series data. This indicates moderate successional patterns over the year, beginning with a single ecostate in April, transitioning to a combination of two states for the summer, then a third state for the late autumn and early winter, before finally transitioning back to the original state in the late winter. This is consistent with the broader seasonality observed in this time series, as suggested in the original paper presenting the entire six-year time series from which the data are excerpted. Without additional context – more intensive temporal sampling, another spatial location, or envionmental covariates – it is unclear if the painting reveals genuine seasonal shifts or other heterogeneity that simply coincides with this pattern, although reports on the larger data set indicate the latter.

**Figure 4:**
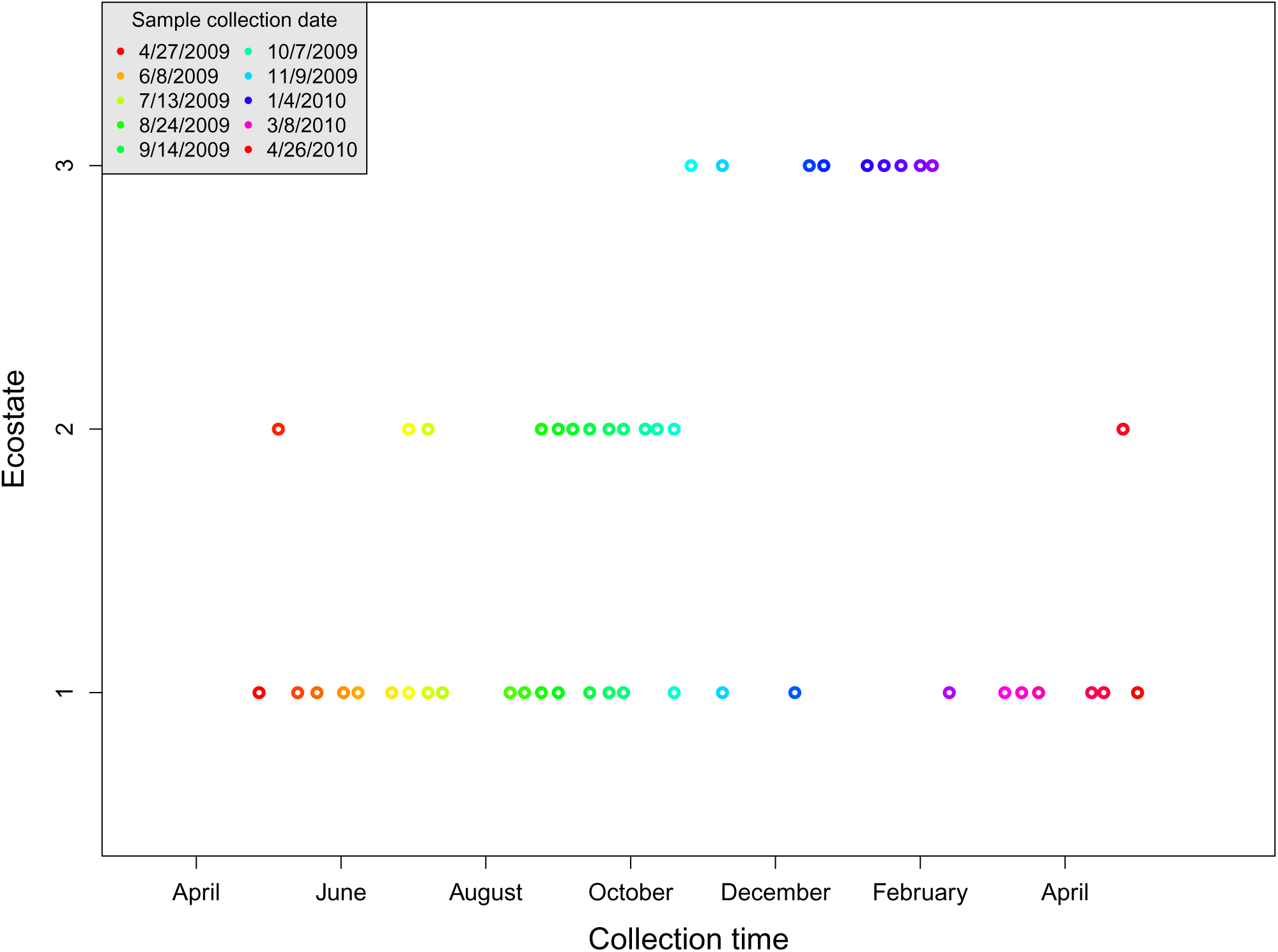
Presentation of ecostates for L4 Western Channel time series. Both x-axis and color denotes time in order to distinguish paired samples. Ecostate shown on y-axis.

**Figure 5:**
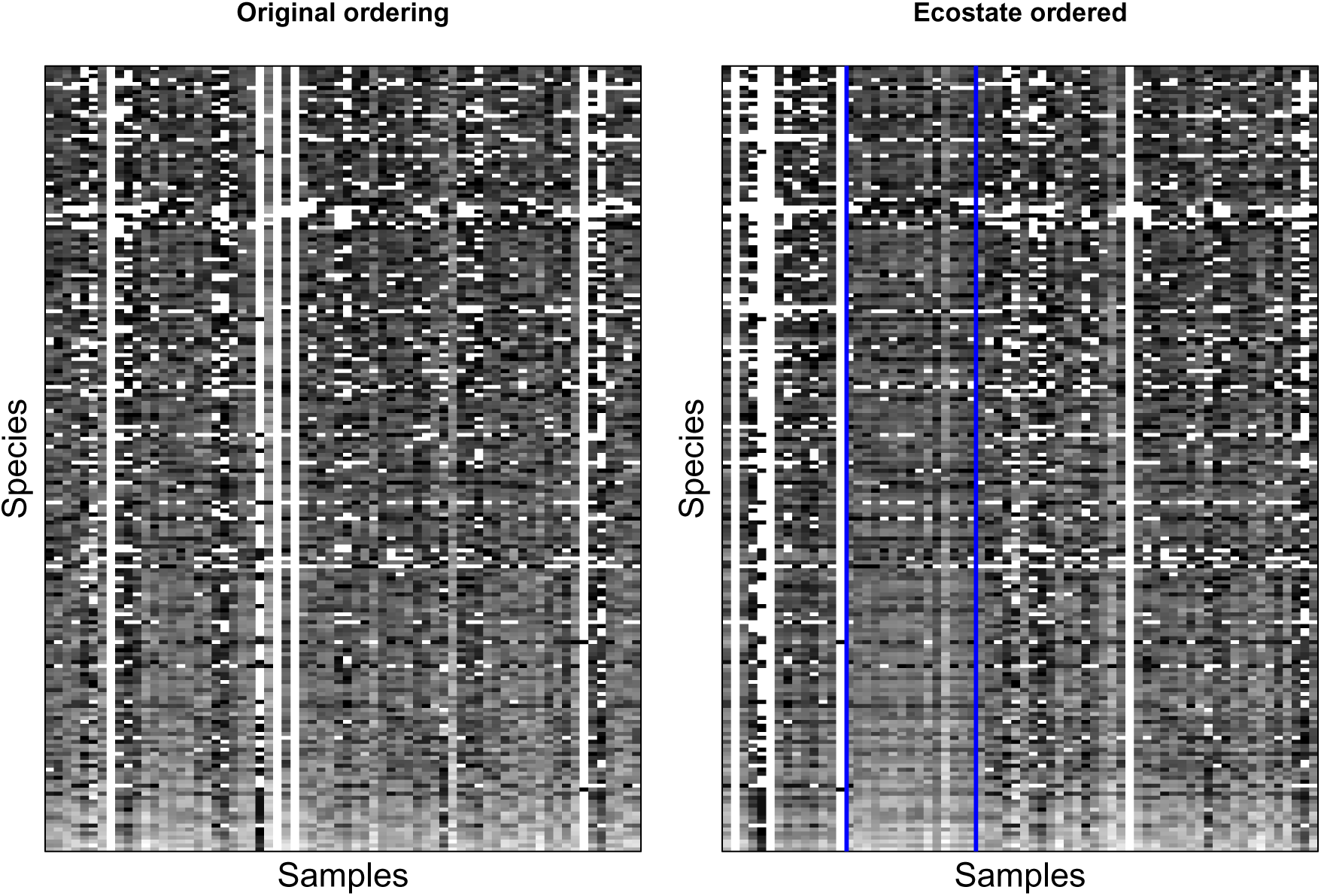
L4 time series data set log-abundance arranged as in initial data set (left) and by maximum-likelihood ecostate (right).

Figure 6 summarizes the PPD analysis for the L4 time series. The *p*-value for each species within each sample shows no systematic issues, except for poor performance across six low count samples (horizontal bands). The model appears to precisely capture mean frequency across samples for nearly all species, while systematically overestimating the amount of variation in counts across samples. The number of zeros within a sample are generally well-modeled, though a significant fraction of samples have overestimates of these numbers. As to the pairwise correlations across species, the model performs generally well for most pairs, although the model overestimates the correlation for a distinct fraction of most frequent species. For moderately frequent and infrequent species, the model shows excellent agreement with the data. In this context, where a single location is observed at moderate temporal resoltion, the iDMM appears to perform very well.

**Figure 6:**
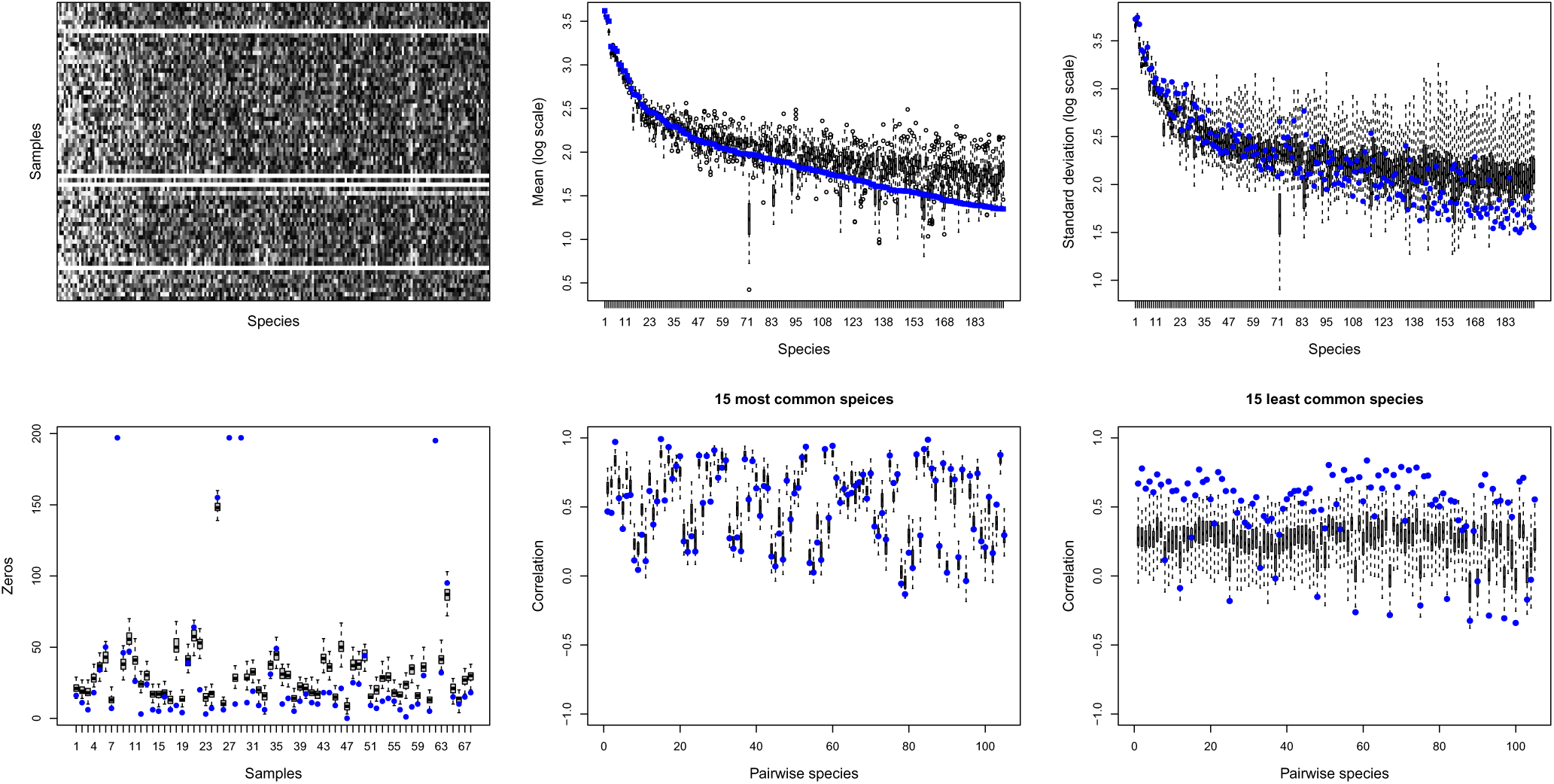
PPD summary for L4 amplicon time series. Note the low count samples contributing the white horizontal bands in the upper left hand panel.

### Gulf of Maine copepod time series

Figure 7 displays the ecological painting for the GoM copepod time series, with each panel organized from left to right by month and bottom to top by distance from Boston, Massachusetts. The painting shows strong seasonal and spatial trends, as frequently noted in the literature (Pershing et al., 2005; Record et al., 2010; Stamieszkin et al., 2015). Both the mid-summer months (Jun.-Aug.) and mid-winter months (Nov.-Feb.) exhibit substantially less spatial variation than other months. The ecostates associated with these relatively quiescent periods show a single or small number of dominant species, with relatively high variation (see Figure 10). The transitional ecostates between these periods exhibit more complex taxon composition.

**Figure 7:**
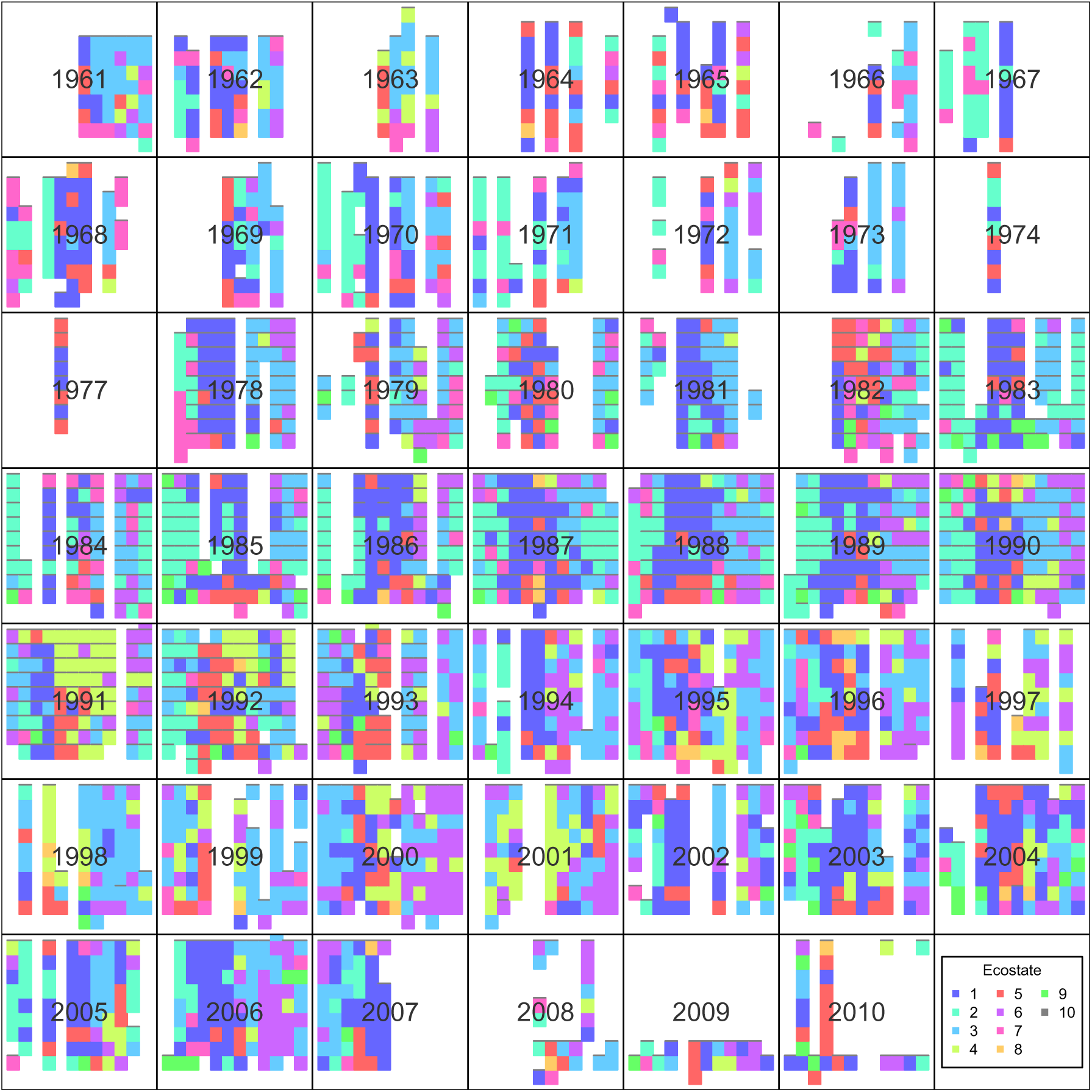
Ecological painting of GoM copepod data set, organized by year. Each panel corresponds to a year, with samples arranged spatially with Boston, USA at the bottom and Yarmouth, Canada at the top and time on the horizontal axis.

**Figure 8:**
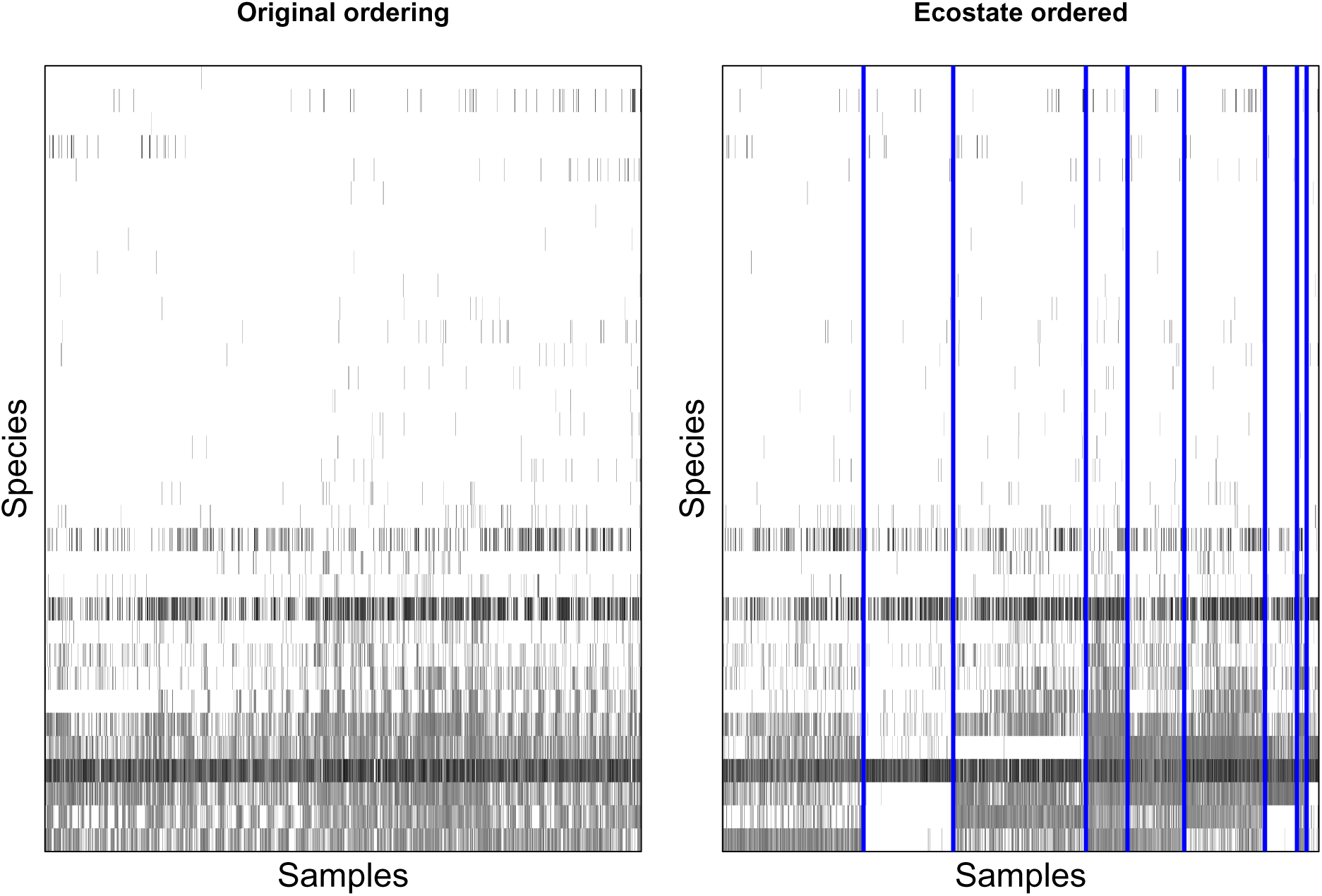
Copepod data set log-abundance arranged as in initial data set (left) and by maximum-likelihood ecostate (right).

**Figure 9:**
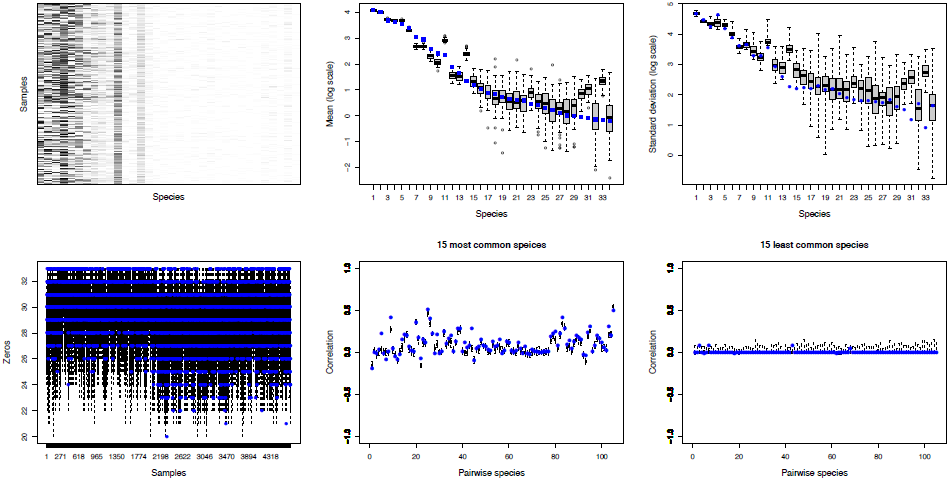
PPD summary for GOM copepod data set. Strong agreement for nearly all aggregate metrics is opposed to poor performance by less frequent species within samples. Note the correspondence between high abundance and low *p*-values in the upper left panel.

**Figure 10:**
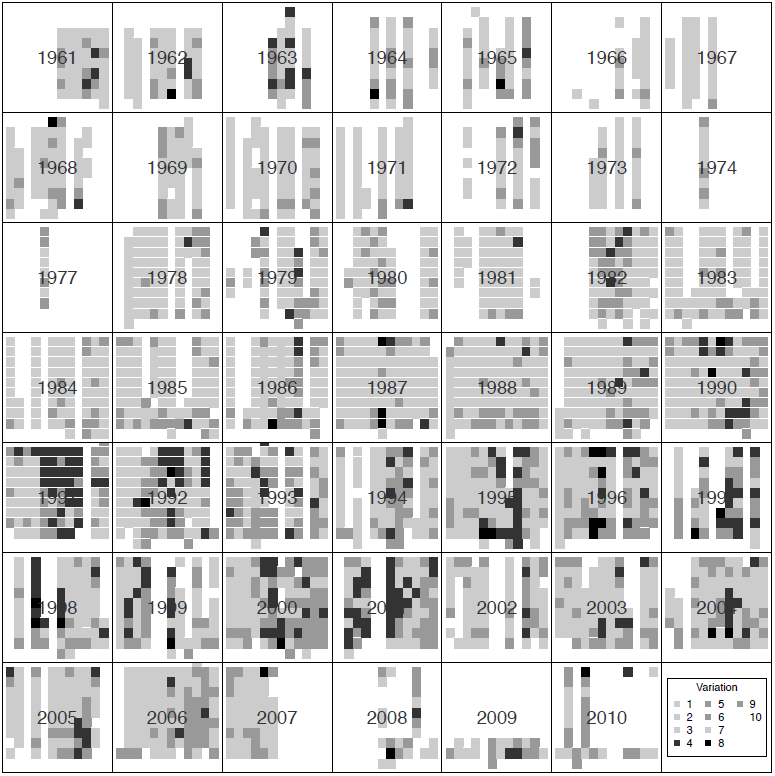
Ecological painting with inverse variance parameter for each ecostate substituted for ecostate coloring. Black indicates high variance; white indicates low variance.

**Figure 11:**
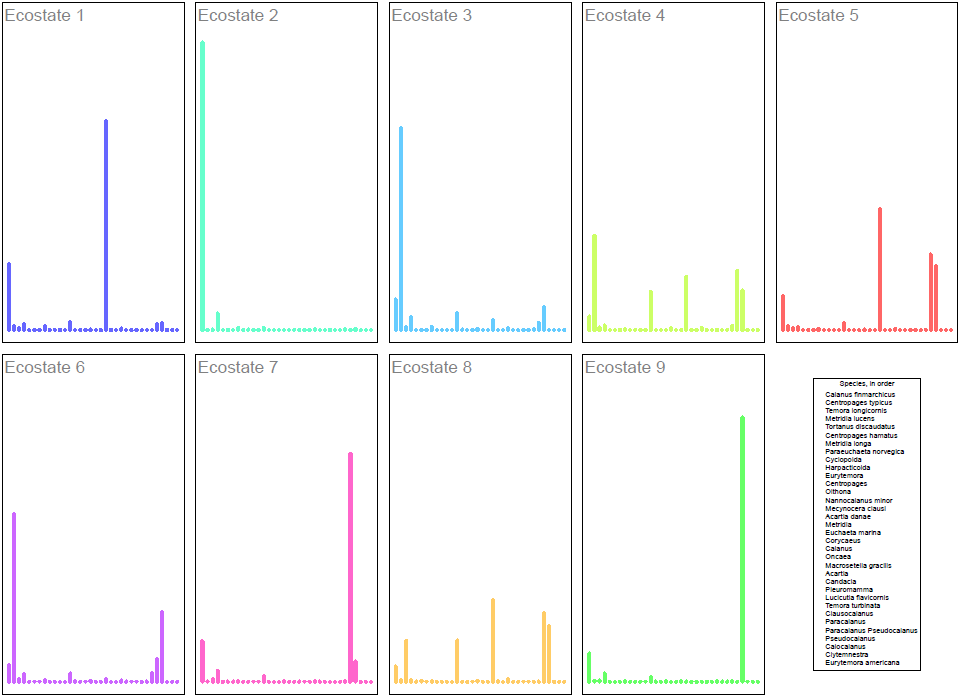
Expected species frequency for each ecostate from maximum likelihood iteration of GoM data set. Colors correspond to Figure 7.

The painting also highlights the dramatic shift in species that occurred from 1990-2001 relative to the years before and after (Record et al., 2010; Johnson et al., 2011). In the years outside the shift, the winter-summer cycle is fairly stable, with only a short transition period associated with other ecostates. This transition is largely dominated by changes in the frequency of *Calanus finmarchicus*, a crucial species in the north Atlantic food web. During this shift, ecological stability diminishes, with transitions associated with more defuse taxa distributions within the present ecostates. During this period, neighboring samples in time and space are less likely to share an ecostate and novel, highly complex ecostates are more common, suggesting ecological distress such as the changes in GoM stratification driven by increased transport of water from the Arctic described in (Greene et al., 2012).

Figure 9 shows the inferred PPD for the dataset. Considering the species/samples values, the performance for highly abundant species is generally good, with poor performance for all other taxa (note the correspondence between *p*-value and mean; upper right and center panels). However, we see strong performance for the mean, standard deviation, zeros, and pairwise correlation. This is consistent with an ecosystem largely determined by a small number of highly abundant species that are likely not exchangeable (in an ecological sense) with other taxa, contrary to the the UNTB. Consequently, the model fits those species well but captures poorly the remaining species. In context, the GoM is at the southern boundary of the range of these taxa. The non-interchangeability revealed here communicates an important vulnerability of this ecosystem to warming.

## Discussion

This work investigates the performance of an infinite dimensional version of the DMM for the ecological count data. It shows how this approach – like other implementations of the DMM – can be used to organize and interpret underlying sample dyanmics with a straight-forward visualization while guarding against overinterpretation by examining the posterior predictive fit. While the model effectively distills many of the important patterns within the data inferred via other non-model-based approaches, such as PCA or MDS, it also shows that some caution should be exercised in these analyses and suggests certain practices may improve the reliability of field investigations.

Posterior checks, especially PPDs, can be an important bulwark in ensuring that these models are treated with appropriate skepticism (Beaumont, 2010). For the DMMs, this requirement is not particularly burdensome as PPDs are easy to simulate and can be interpreted using familar *p*-values as a metric. Appropriately used, these measures give researchers and their readers increased assurance in the reported conclusions. We encountered several issues in the analysis of the datasets above that suggest useful guidance for field researchers. We organize these suggestions below under the following headings: identifying problems with samples; relevant scales of ecological fluctuation; niches as ecological outgroups; and departures from niche neutrality.

### Identifying problems with samples

In our examples, PPDs consistently identify problematic samples from the iDMM’s perspective that may suggest experimental issues. In the cases we present, sample-specific deviations from model expectations may be due to low sample counts, mislabeled samples, unusual taxa present, or departures from the iDMM’s assumptions. PPDs for the iDMM provide a convenient tool for identifying these samples, showing low *p*-value ‘bands’ (bright red) that are indicative of poor performance for a sample across a subset of taxa. In the L4 example, bands across all species indicated insufficient counts for reliable inference within certain samples. In the MPHM painting, the tongue ecostate is found in the feces habitat, a likely mislabeling. The ecological painting provides an easy means to identify these questionable samples.

### Relevant scales of ecological fluctuation

By ecological fluctuation, we mean the changes in environmental conditions that precipitate shifts in taxon composition and consequently ecostate. The easiest scale of ecological fluctution to analyze is the case-control study. As highlighted in their analysis of twin obsesity (Holmes et al., 2012), the association between iDMM components and case/control status yields a straight-forward statistical approach. A similar approach was recently used to analyze the microbiomes of human cancers (Nakatsu et al., 2015). However, ecologists are often interested in more sampling across more heterogeneous patterns of spatial, temporal or environmental change. While there is some guidance about the level of read-depth required for DM inference (La Rosa et al., 2012), there is little guidance available for dealing with ecostate variation.

The analysis of time series datasets here makes it clear that ecologists should prepare a sampling design of sufficient resolution to capture ecostate transitions. In practice, this means sampling at a temporal resolution substantially higher than the expected scale of the transition itself. For instance, the MPHM data is compelling in large part because it shows the relative constancy of species abundance in time and space. However, this consistency is revealed precisely because the sampling occurs at a time interval substantially more frequent than the scale of ecostate variation. Similarly, the absence of intensive sampling renders the conclusion of seasonality in the L4 time series uncertain. A seasonal transition that would be anticipated for a temperate marine microbiome is on the order of 90 days (a single season). A minimum sampling regime should then be an order of magnitude less than that, or less than 9 days, about 60% of the two week average interval for the L4 study. In other cases, such as spatial or environmental gradients, a similar rule of thumb can be employed.

### Niches as ecological outgroups

The specific examples we consider here strongly suggest that researchers consider using ‘outgroup’ sampling as important tool for contextualizing their analyses. The term outgroup in phylogenetic analysis refers to a species included in the collection for the purpose of parsing the relatedness among the study population (Nixon and Carpenter, 1993). In the context of ecological data, we mean a sample that has some related properties to the main study target but is sufficiently dissimilar to provide a backdrop for understanding the degree of variation in the study population. An outgroup does not need to fall entirely outside the study domain. For instance, in the MPHM, the separate niches (feces, saliva, skin) provide simultaneous outgroups for each other, allowing more confident interpretation. In the copepod data set, the extensive temporal and spatial sampling that the study provides is akin to an outgroup, though a set of off-shelf samples would be better. The analysis of the L4 time series suffers from the lack of an outgroup, leaving the apparent seasonality in doubt (though reports from the complete, six year dataset indicate the robustness of this observation).

### Departures from the niche neutrality

To some researchers, departures from the DMM model assumptions mean that the UNTB and analysis based on the DMM can be abandoned immediately (Dornelas et al., 2006). For researchers content to still use this analysis, these departures may be useful: as the data depart from model expectations, the structure of these deviations can reveal important aspects of the ecology. In the GoM copepod data set, the domination by *C. finmarchicus* represents a departure from underlying model assumptions as it cannot be ‘swapped out’ for any other copepod species (i.e. it is not ecologically equivalent).

This is clearly shown in the PPDs for each species: the model performs well for highly abundant taxa like *C. finmarchicus* but poorly for all other taxa. This is consistent with the common hypothesis that *C. finmarchicus* is irreplaceable in the GoM. This species is at the southern boundary of its range in the GoM, implying a high vulnerability of the system to warming. It also shows how a departure from the DMM model assumptions and hence the UNTB can reveal critical characteristics of ecosystem function. We understand that these departures are qualitative (‘bad’ fit with the PPDs) rather than quantitative. An important avenue for future research with the DMM is to develop precise statistical tests for departures from model expectations with ecological understandings of their consequence (Harris et al., 2014).

## Conclusions

The DMM and its variants provide powerful tools for understanding ecological count data. However, these approaches possess a number of weaknesses that could be addressed in the next generation of models. Most obviously, these approaches do not consider the total number of model counts within a sample, a critical indicator of ecosystem function. Researchers might naturally account for this by excluding exceptionally high‐ or low-count samples, though this would be better addressed by including this variation at the level of the model, as is possible with a negative binomial process (Zhou and Carin, 2015). The iDMM also does not account for correlation across samples, as could be done using Gaussian process priors or hidden Markov models to ‘borrow strength’ across samples.

Finally, we do not believe that using the iDMM requires one to take a strong position on the UNTB: the underlying ecostates can be treated as phenomenological clusters, rather than theoretically-precise metacommunity structures, and still provide analytic utility, as evidenced particularly well in the copepod data set. As the use of the coalescent model in phylogenetics is not substantially diminished by the frequent observation of violations of its assumptions, the approach of the iDMM and similar models promise the possibility of a connection to the broader UNTB framework while often giving a practical means of interpreting datasets independent of its operation.

## 1 Acknowledgements

The authors declare no competing interests. We gratefully thank Ana Lagunez for careful editing of the manuscript.

